# The role of dormant propagule banks in shaping eco-evolutionary dynamics of community assembly

**DOI:** 10.1101/2024.11.03.621703

**Authors:** Jian-Hao Lin, Bo-Ping Han, Mark C. Urban, Luc De Meester

## Abstract

The metacommunity framework provides a theoretical framework for understanding how dispersal alters species distribution patterns at different spatial scales. Dormancy is a strategy that decreases the extinction risk and can be viewed as dispersal over time, shaping the assembly dynamics of populations and communities. However, dormancy has only recently been recognized as a potentially important mediator of metacommunity-level processes. Here, we extend an individual-based model of metacommunity dynamics by adding propagule banks that store dormant propagules that can hatch in subsequent growing seasons. By systematically varying the propagule bank size in the model, we investigated how dormant propagule bank size influenced the eco-evolutionary dynamics of community assembly in populations of both asexual and sexual species at three dispersal levels (isolated communities and weakly and strongly connected communities). Our simulations show that as the propagule bank size increases, 1) the rate of evolution decreases in asexual species, whereas the rate of evolution increases in sexual species; 2) species sorting becomes less important while evolution-mediated zpriority effects (monopolization) make a more significant contribution; and 3) α-diversity decreases and β-diversity increases. We also found that patches with well-established propagule banks contributed more to the regional species pool than patches with less well-developed propagule banks. Dispersal asymmetries also emerge as patches with a well-established bank tend to reduce the invasion success of new arrivals while also providing more emigrants with diverse genetic variation to the other patches, fostering regional monopolization. To this end, we regard the banks in a metacommunity as a hidden state of a local community, a potential gene pool, and a latent part of a regional species pool.

## Introduction

A key goal of ecology is to understand both local and regional community patterns (Ricklefs 1987, Leibold et al. 2004, Cornell & Harrison 2014, Urban et al. 2020). In the classic metacommunity framework, populations and communities at a regional scale are linked by dispersal, and all individuals of all species in the local patches of a metacommunity collectively constitute the regional species pool, whereas the metacommunity is also affected by the mainland as an external species pool (Fukami 2005, 2015). Local community assembly results from the dispersal of individuals, drift, and selection by the environment (Vallend 2010). More specifically, the internal structure of a metacommunity can also influence the resulting community, with differences among species in terms of their niche and dispersal traits, their relative abundances, the connectivity of a patch, and its location (Leibold et al 2022a). However, in addition to purely ecological dynamics, there is the potential for eco-evolutionary interactions to impact local community assembly (Leibold et al 2022b). Eco-evolutionary feedback, which has the potential to strongly influence the structure of evolving metacommunities (Urban & Skelly 2006, Urban et al 2008), is the so-called evolution-mediated priority effect (or monopolization; De Meester et al 2022, Loeuille & Leibold 2008, De Meester et al 2016). Evolution-mediated priority effects occur when early colonists undergo adaptive evolution, increasing their fitness, thereby reducing the establishment of later colonists and finally monopolizing a community (Urban and Skelly 2006, Fukami 2015, De Meester et al. 2016). The monopolization effect can alter community assembly dynamics and produce contingent patterns that deviate from expectations based on species sorting and are more similar to neutral-like dynamics (De Meester et al. 2002, Loeuille and Leibold 2008, Urban and De Meester 2009, Waters et al. 2013, Vanoverbeke et al. 2016, Leibold et al., 2019).

Dormancy is a common feature of many taxonomic groups, including plants and algae, aquatic animals such as bryozoans, cladocerans, copepods, and rotifers, and terrestrial animals such as nematodes, some insects, and even some Cyprinodontidae fish (Templeton and Levin 1979, Perry 1989, Bailey 1972, Koštál 2006). Seeds of many annual plant species can remain dormant for numerous years, so the abundance of annual plants does not merely depend on the last year’s seeds but the germination of the accumulated seed banks from preceding years as well (Templeton and Levin 1979). Similarly, freshwater zooplankton propagule banks are composed of dormant eggs that accumulate in lake sediments and are often viable for long periods (Hairston et al. 1995, Hairston 1996a, Cáceres 1997a, Geerts et al. 2019). Although dormant propagule banks have a high potential to influence a metacommunity assembly, both classical and evolving metacommunity theories generally ignore dormancy (Wisnoski et al. 2019).

Dormancy and germination of propagules in banks can alter the ecological and evolutionary dynamics of the local population and community assembly (Pavone and Reader 1982, Dalling et al. 1998, De Meester et al. 2002, 2016). As the propagule bank size increases, it can significantly influence the dynamics of local populations (Mistro et al. 2005, Wisnoski and Shoemaker 2021). For species living in fluctuating environments, propagule banks can also affect the rate and direction of population and community responses to environmental changes (Chesson 1994, Hairston 1996a, Hairston and Cáceres 1996). Propagule banks can buffer fluctuations in the population size of active populations (Warner and Chesson 1985, Hairston et al. 1996b, Cáceres 1997b) and constrain later arriving genotypes or species that initially lack this propagule bank (De Meester et al. 2002, Holyoak et al. 2020), which is the storage effect (Templeton & Levin 1979, Warner & Chesson 1985, Hedrick 1995). Propagule bank size is likely a crucial parameter determining the effects of dormancy on population and community dynamics. Propagule bank size of a given population is expected to vary with the species’ traits such as size and shape of the dormant stage, life history strategies determining when and how many dormant propagules are produced (Pavone and Reader 1982, Thompson et al. 1993, Hulme 1998, Aikio et al. 2002, Brown and Venable 1991), as well as species interactions within the community and local environmental conditions that determine local abundances (Holyoak et al. 2020; Wisnoski and Shoemaker 2021).

Propagule banks can be strongly affected by and influence the metacommunity structure. Locally, bidirectional feedback exists between the size and dynamics of propagule banks (e.g., storage effect, Warner & Chesson 1985) and local population and community dynamics (e.g., selection, species interactions, Vallend 2010). Regionally, banks can affect a regional species pool, and vice versa, by providing potential immigrants with new or empty habitats (Zobel et al. 1998, Harrison and Cornell 2008, Lessard et al. 2012, Fukami et al. 2004, 2015, Zobel et al. 2011, Okamura and Freeland 2002, Wisnoski et al. 2019). The dynamic dependence between a metacommunity and its regional species pool has been largely ignored in the classic metacommunity theory, leading to a relatively static state of the species pool (Mittelbach & Schemske 2015). Dormancy has recently been proposed as an important metacommunity-level process (Wisnoski et al. 2019, Wisnoski and Shoemaker 2021, Holyoak et al. 2020). Dormancy and dispersal represent similar strategies used by species to reduce extinction risk in variable environments (Buoro and Carlson 2014), both modifying metacommunity dynamics. A large propagule bank in a patch is expected to compensate for local population losses and extinctions, reduce dispersal success from other patches, and increase the number of potential migrants to other patches (Wisnoski et al., 2019). In addition, propagule banks not only enhance species persistence but also their genetic diversity. The effect of propagule banks on genetic diversity (Hedrick 1995) and the evolutionary rate of traits under fluctuating selection (Hairston 1998) implies that propagule banks also influence genetic responses to environmental gradients and species interactions. Therefore, dormant propagule banks can strongly influence eco-evolutionary dynamics in metacommunities. Although the original monopolization hypothesis emphasizes the importance of dormant propagule banks in fostering evolution-mediated priority effects (De Meester et al 2002), nevertheless, the importance of dormant propagule banks has not been incorporated into theoretical models.

In the present study, we modified an existing individual-based model that incorporates evolution in metacommunities (Vanoverbeke et al. 2016) to include propagule banks and propagule dispersal. We aimed to better understand how and to what extent propagule bank size and dispersal in interaction with eco-evolutionary feedback influence the structure of metacommunities. We focus on how propagule bank size and dispersal level affect: (1) the dynamics of evolution of local pioneer populations, (2) the dynamics of early colonizers and establishment success of later immigrants, (3) the relative importance of assembly mechanisms across the metacommunity, and (4) α, β and γ-diversity of communities at habitat-, patch- and regional scales.

## Models and methods

We adopted the structure of a previously developed metacommunity model (Urban and De Meester, 2009, Vanoverbeke et al. 2016), but additionally assumed that each patch in the metacommunity is connected by limited passive dispersal from a local dormant propagule bank. Our model is an integrated model that combines both a classic island-mainland model and a metacommunity model (MacArthur & Wilson 1967, Leibold et al. 2004, Fukami 2005, 2015). Similar to the island-mainland model, the mainland acts as a large external species pool that provides continuous propagule rains to the initially empty metacommunity throughout the simulation. The initially empty patches were under the propagule rains of four species from the mainland, with two traits corresponding to two types of environments in the landscape. From the initial empty patches, the metacommunity started with propagule rains, and then the species could also disperse across the metacommunity, gradually establishing an internal species pool, which is the metacommunity itself. Finally, metacommunity dynamics are under colonization by both the mainland and internal species pools.

We established a hierarchically structured metacommunity comprising four regions. Each region contained four patches and each patch had four habitat types (Fig. 1a). We created two hierarchically structured environments: at the metacommunity level, the four environmentally different regions (denoted as *RE in the remainder of the text*, i.e., regional environmental value) had mean *RE* value of 0.2, 0.4, 0.6, and 0.8, denoted as green in Fig. 1a; at the patch level, we set four environmentally different habitats (denoted as *HE* in the remainder of the text, i.e., habitat environmental value), and the mean of the *HE* values in each habitat was 0.2, 0.4, 0.6, and 0.8, denoted as blue in Fig. 1a. Note that *RE* and *HE* are two different environmental axes. In the mainland. the mean values of the two traits of the four species, which correspond to *RE* and *HE* respectively, are (0.2, 0.2), (0.4, 0.4), (0.6, 0.6), and (0.8, 0.8). All the species had equal dispersal abilities. We ran each model for 5000 time-steps to reach quasi-equilibrium, which was confirmed by further simulations up to 20000 time-steps (Fig. 4A in Appendix 4, Videos in Appendix 6).

**Fig. 1.**
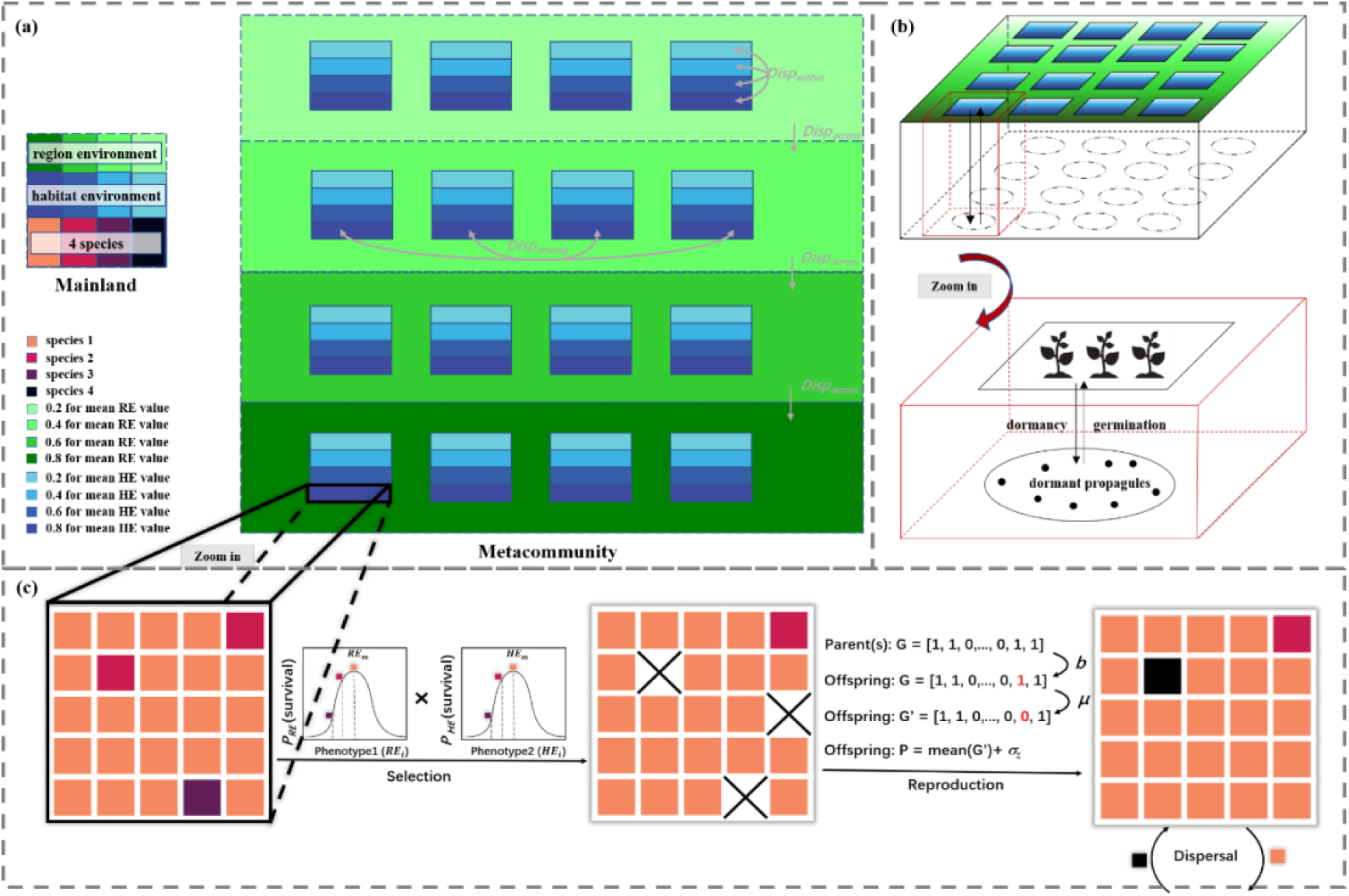
The model extends the one by Vanoverbeke et al. (2016) by adding propagule banks. The metacommunity includes 3 spatial scales: habitats, patches, and regions. (a) The habitat environment values (HE values) vary within patches (with mean values of 0.2, 0.4, 0.6, 0.8; blue) and the regional environmental values (RE) vary across regions in the metacommunity (with mean values of 0.2, 0.4, 0.6, 0.8; green; HE and RE differ along different environmental axes). Individuals undergo death, reproduction, mutation, dispersal, and dormancy processes in the simulation. At the start of the simulation, all the patches in the metacommunity are empty and the initially-occupied mainland provides a source for the propagules of four species (denoted as black, purple, red, and orange). In the metacommunity, offspring can disperse within a patch, among patches and across the region with dispersal rates, denoted as *m*_*within*_, *m*_*among*_ and *m*_*across*_ respectively. (b) A conceptual graph of the relationship between local patches and the resting propagule bank. In each time step, the existing offspring can be stored in the bank via the dormancy process and the dormant propagules can occupy empty sites by germination. (c) Following mortality selection, individuals can reproduce asexually or sexually and offspring genotypes are based on that of their parent(s). Mutation occurs during reproduction when the bi-allelic gene changes value from 0 to 1 or vice versa, with a mutation rate μ. We model *L = 40* bi-allelic genes and assume that they determine trait values additively. We calculate the two phenotypes (denoted as *RE*_*i*_, *HE*_*i*_, corresponding to the RE-related phenotype value or HE-related phenotype value of an offspring individual, respectively) as the means of the first or second set of 20 bi-allelic genes plus a Gaussian random variance (*σ_z_ = 0 025*) simulating an environmental influence on species traits.

At the start of the simulation, we created an empty metacommunity, initializing the specific microenvironment values of all microsites. Each patch consisted of 100 micro-sites. Each microsite had two specific microenvironment values of *RE* and *HE* (named RE microenvironment and HE microenvironment, denoted as *RE*_*m*_ and *HE*_*m*_, respectively). In each microsite, the *RE* micro-environment value (*RE*_*m*_) is equal to the mean value of *RE* (according to which region is the microsite located) plus a stochastic Gaussian variable (σ_*RE*_ = 0.025), and the *HE* micro-environment value (*HE*_*m*_) is also equal to the mean value of *HE* (according to which habitat is the microsite located in) plus a stochastic Gaussian variable (σ_*HE*_ = 0.025). We then created an initially occupied mainland with four species (denoted *sp1*, *sp2*, *sp3,* and *sp4* as *species identifier* below). Each species has two traits, the first of which is a RE-related trait and the second is a *HE*-related trait. The mean *RE*-related- and *HE*-related trait values for each species were 0.2, 0.2 for *sp1*, 0.4,0.4 for *sp2*, 0.6, 0.6 for *sp3* and 0.8, 0.8 for *sp4,* respectively. In the initialization of specific individuals on the mainland, the phenotype is genetically based, and we assumed *L* = 40 bi-allelic additive genes coded as 1s and 0s, where the first and second 20 genes control the RE-related phenotype of an individual (*RE*_*i*_) and the *HE*-related phenotype of an individual (*HE*_*i*_), respectively. First, we set the genotypes of an individual such that the mean value of the genotypes was equal to the mean trait value. Then, the RE-related phenotype of an individual (denoted as *RE*_*i*_ below) or HE-related phenotype of an individual (denoted as *HE*_*i*_ below) is equal to the mean value of RE-related trait values (*RE*) or the mean value of HE-related trait values (*HE*) plus a stochastic Gaussian variable (σ_*RE*_ = σ_*HE*_ = 0.045), representing the standing genetic variation (Barrett and Schluter 2008).

We applied a lottery model in which each microsite can only be occupied by one individual and only the empty microsites can be colonized, with each individual undergoing selection. Following Vanoverbeke et al. (2016), we calculated the survival rate for a given individual in each microsite as a Gaussian distribution:

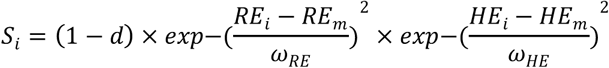

where *d* = 0.1 is the baseline mortality rate as the disturbance, ω_*RE*_ = ω_*HE*_ = 0.5 is the width of stabilizing selection, *RE*_*i*_, *HE*_*i*_ is the RE-related phenotype value or HE-related phenotype value of an individual and *RE*_*m*_, *HE*_*m*_ is the RE-related micro-environment value or the HE-related micro-environment value in a specific microsite. At each time step, all individuals in the metacommunity underwent natural selection. We generated a random variable (*u*) ranging from 0 to 1 that follows a uniform distribution, and if *u > S*_*i*_, the individual undergoes mortality and the microsite becomes empty.

After selection, surviving individuals reproduced asexually or sexually. Sexual individuals can mate only with the same *species*. Each value of the biallelic genes was inherited from the female or male parents with the same probability (i.e., 50 %). In asexual individuals, the genotype of the offspring was identical to that of the parent. If a mutation occurs, the bi-allelic gene values change from 1 to 0 (vice versa), with the mutation rate, μ = 10^−4^ The *RE*-related phenotype value (*RE*_*i*_) and HE-related phenotype value (*HE*_*i*_) of a specific offspring are calculated as the mean of the first 20 or second 20 additive gene values plus a non-genetic phenotypic variation (σ_*z*_ = 0.025). The total number of offspring (*I*) across the metacommunity at each time step is

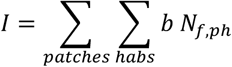

where b is the birth rate depending on the reproductive strategy (i.e., 0.5, asexual, 1 sexual) and *N*_*f,ph*_is the number of females in habitat *h*, patch *p*. Note that in the asexual mode, all the individuals were female, and in the sexual mode, half of the individuals were expected to be female. *patches=16* is the number of patches in the metacommunity and *habs = 4* is the number of habitats within a patch.

In modeling the dispersal process, our model assumes that only dormant propagules can disperse between habitats, patches, or regions and that microsites can be occupied by either local offspring or migrant dormant propagules. We assumed three dispersal pathways: dispersal across regions, dispersal among patches within a region, and dispersal within a patch, which are controlled by the dispersal rates denoted as *m*_*across*_, *m*_*among*_, and *m*_*within*_. In this spatially explicit model, dispersal across regions is long-distance dispersal, dispersal among patches within a region is medium-distance dispersal, and dispersal within a patch is short-distance dispersal. Therefore, we set the dispersal rates such that *m*_*across*_ < *m*_*among*_ < *m*_*within*_. The expected number of emigrants leaving *patch p* by dispersal *pathways q* per time step, *Disp*_*p,q*_, is calculated as

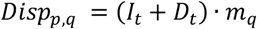

where *I*_*t*_ is the total number of non-dormant offspring in the patch; *D*_*t*_is the total number of dormant propagules in the bank at *time-step t* (for the definition and calculation of *D*_*t*_, see below); and *m*_*q*_ is the dispersal rate, which depends on the dispersal pathways, where q denotes *across regions, among patches within regions* or *within a patch*. In the next time step, the immigrant species will compete with the local species (existing offspring and germinated dormant propagules) for empty microsites by random sampling.

Finally, in the dormancy process, the propagule bank stores all the newborn propagules from the local community in the bank; that is, the banks can provide dormant offspring for passive dispersal and germination (Okamura and Freeland 2002). At the end of each time step, the existing active offspring enter dormancy in a local propagule bank, which contains both dormant offspring from previous time steps and new dormant offspring from the current time step. In the model, we assumed that the propagule bank size was dynamic but constrained by a fixed carrying capacity. To achieve this, propagules were randomly eliminated when the maximum was exceeded. This constant carrying capacity for dormant egg banks is somewhat artificial; dormant propagule banks for any community also suffer mortality (e.g., propagule predation), and keeping the dormant bank at a given size allowed us to systematically analyze the impact of dormant propagule banks on metacommunity dynamics. The number of offspring in the propagule bank at each time step was as follows:

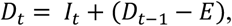

where *D*_*t*_ is the total number of offspring in the bank at time step t and is equal to the number of newborn offspring at time step t (*I*_*t*_) plus the number of resting offspring inherited from the pool at time step t-1 (*D*_*t*−1_) minus the number of randomly removed dormant offspring in the bank (*E*; *E* is adjusted such that it keeps the total number of offspring in the bank within its carrying capacity). In other word, we note that if *I*_*t*_+*D*_*t*−1_ ≤ *the propagule bank capacity*, then no individuals will be eliminated i.e., E = 0 (Fig. 2). All parameters and values used in the model are listed in Table 3S in Appendix 7.

**Fig. 2.**
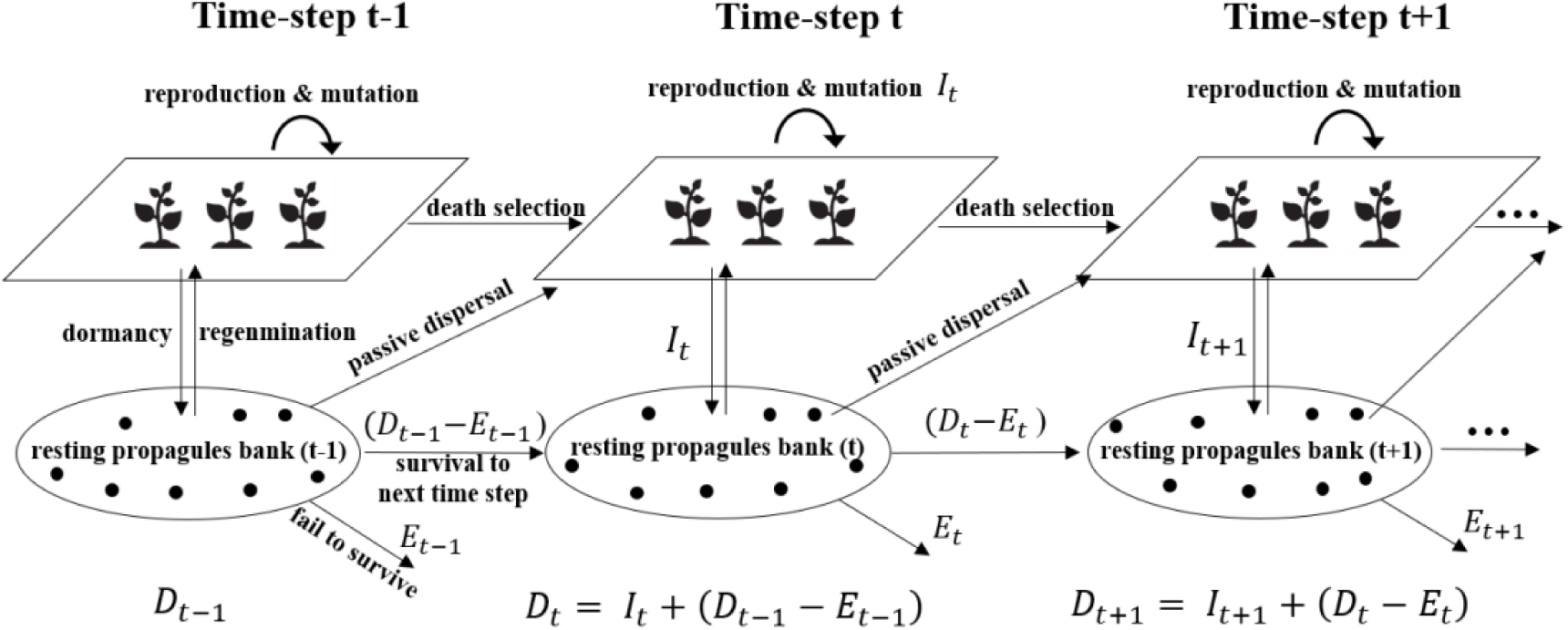
The relationship between the metacommunity dynamics (top panels) and the dormant propagule bank dynamics (lower panels) at time step t-1, t, and t+1. Individuals in the model undergo mortality, selection, reproduction and mutation, dormancy in the bank, dispersal, and dormant propagule elimination processes. At time-step t, after passive dispersal, individuals undergo mortality selection followed by reproduction and mutation. The empty sites in the metacommunity can be occupied by individuals through random sampling with replacement from the banks via re-germination, the existing offspring via reproduction or exotic individuals via dispersal. We set a parameter, *‘the dormant propagule bank size’* to control the maximum capacity of the bank. *D*_*t*_., The total number of offspring in the propagule bank at time step t is expected to be all the new-born offspring from time step t (*I*_*t*_) plus the dormant ones remaining in the pool at time step t-1 (*D*_*t*−1_) and minus a proportion of offspring in the bank that is randomly eliminated to keep the bank size within the carrying capacity. Note that the new dormant offspring added into the bank by the dormancy process at time-step t will not be eliminated if *I*_*t*_+*D*_*t*−1_ ≤ *the propagule bank size*, i.e., E = 0.

At the end of the simulation, we measured biodiversity at three spatial scales using the inverse Simpson’s index, *D* = 1/ ∑ *p*_*i*_^2^ (Simpson 1949). Alpha diversity (α_D_) in habitats, patches, and regions is defined as the inverse Simpson’s index at these scales, and gamma diversity (γ_D_) is defined as the inverse Simpson’s index of the global metacommunity. Following Whittaker (1960), beta diversity (β_D_) is defined as the ratio of gamma diversity to alpha diversity: β_D_ = γ_D_/α_D._ The ratio of gamma diversity to alpha diversity in patches indicated the variance in species composition between patches, and the ratio of gamma diversity to alpha diversity in regions indicated the variance in species composition between regions.

Next, we evaluated the variable contributions of individual assembly mechanisms, including the ecological mechanisms of species sorting and mass effects, and eco-evolutionary mechanisms, including the monopolization effect, according to the decision-making table for assemblage mechanisms (after Vanoverbeke et al. 2016, Table 2 in Appendix 5). If a portion of over 90 % of the microsites in the local habitat are occupied by the same species (in this case, α-diversity is 1.25), the species are assumed to dominate the local habitat. The value of 1.25 is used to delimit whether a local habitat is dominated by one single species (diversity < 1.25) or experiences a mass effect (diversity≥1.25). When diversity was < 1.25, a monopolization effect was identified when one species dominated and was not originally pre-adapted to that locality; otherwise, species sorting was performed.

We also estimated the rate of adaptation of a population to the local habitat as the log change in trait values scaled by the pooled standard variation per generation (*haldanes*) (Hendry and Kinnison 1999, Appendix 2 for details). Notably, the generation time for an evolving population is not a constant, but related to the fitness of a population (i.e., the mean survival rate of a population) to the local environment dynamically (Gotelli 2008, see calculation details in Appendix 3).

### Simulation scenarios

To understand how propagule bank size affects a metacommunity, we simulated the model by setting the parameter *‘bank size’*, i.e., the maximum capacity of the dynamic bank size, from 0 to 8000 in steps of 800. We ran the simulation with three dispersal rate levels, that is, isolation, low, intermediate, and high dispersal (values of dispersal scenarios in Table 1), each for 5000 *time-steps* resulting in quasi-equilibrium, which was confirmed with simulations of 20000 time-steps. Simulations were run for both sexual and asexual reproduction scenarios. We ran the model for each set of parameters with *r = 60* replicates and 5000 *time-steps*.

By simulating the dormancy process, we sought to understand: 1) how dormancy affects local population dynamics when other metacommunity processes are excluded, 2) the effect of dormancy when combined with other metacommunity processes, and 3) the effect of dormancy on eco-evolutionary dynamics in an evolving metacommunity. A scenario involving an isolated population was implemented to test the first goal of simulating the microevolution of a local population in the absence of gene flow, source-sink dynamics, and interspecific competition. In this scenario, we modeled only one species and explored how maladapted species can adapt to different local patches when dormant propagule banks of different sizes are presented (see details in Appendix 2). Different dispersal scenarios are required to understand the interactions between dormant propagule banks and other metacommunity processes, whereas allowing for evolution favors us evaluating how dormancy affects eco-evolutionary dynamics in an evolving metacommunity.

We coded the simulation model in Python 3.7 and we used the *mpi4py* library to run all the simulation trials on the High-performance Computing Platform of Jinan University.

## Results

### Local genetic adaptation as a function of the size of the dormant propagule bank

Under simulations with a single species in an isolated habitat, dormancy has a strong impact on the changes in average fitness (quantified as the mean survival rate of the population) over time (Fig. 3), the evolutionary rate quantified as the mean *‘log10 haldanes’* between each 10 generations during the entire simulation time (Fig. 3A in Appendix 2), and the dynamics of population abundances in the entire metacommunity (Fig. 2A in Appendix 2).

**Fig. 3.**
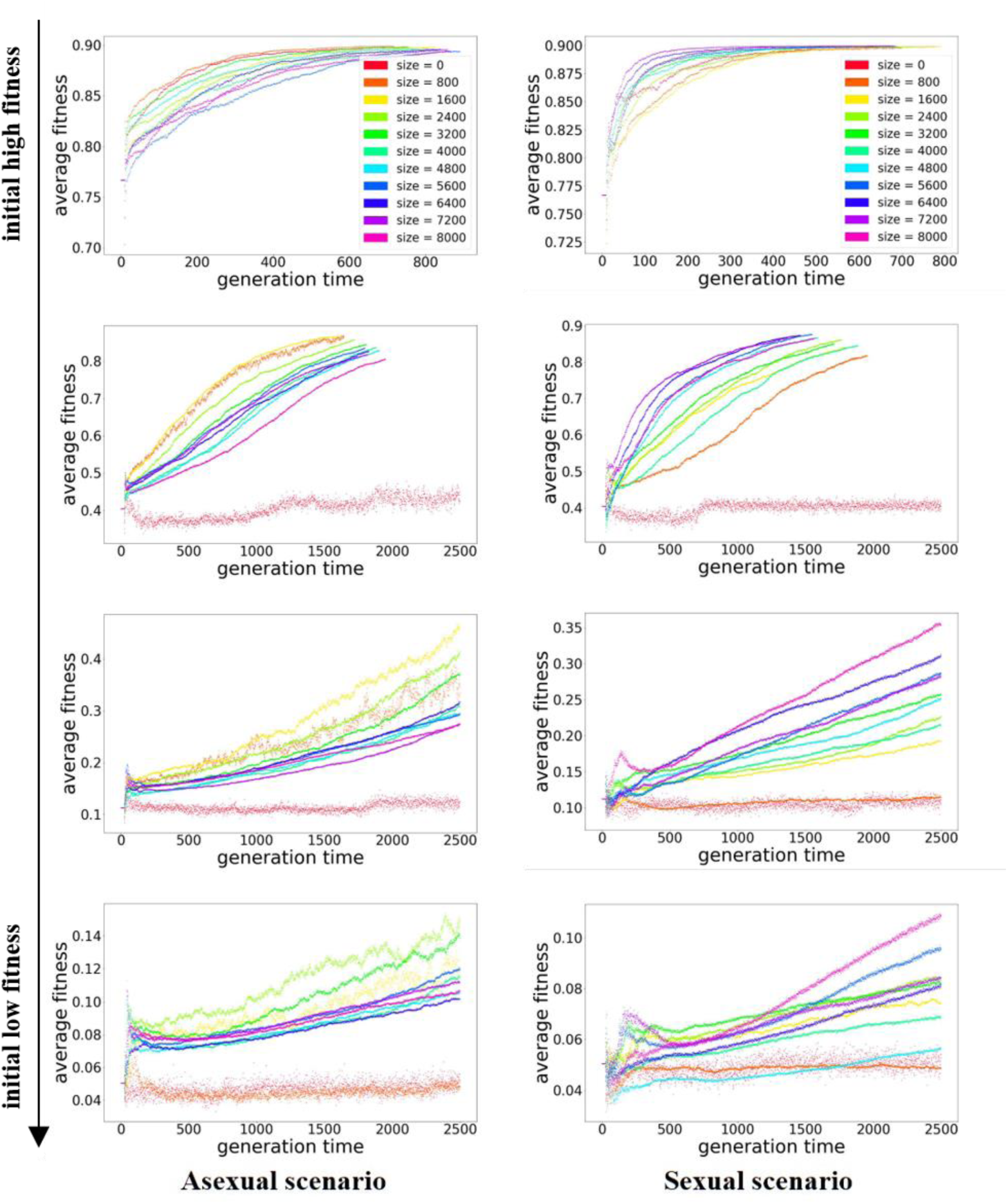
The evolutionary change in fitness of an initially maladapted founding population in the presence of dormant propagule banks of different sizes and under a scenario of asexual and sexual reproduction. The average fitness of the local population is calculated as the mean survival rate based on the match between its traits and the environmental values. We ran the model with 5000 time-steps in total, and the generation time of an evolving population is not a fixed constant but a dynamic one. Usually, the higher fitness a population is, the longer the generation time of the population is, i.e., G = 1 / (1-S) in Appendix 3. Then, we rescaled the x-axis of the plotting from time-steps scales to generation time scales, based on their fitness data. Note that under the condition of initial low fitness and no or small bank, the local population repeatedly underwent colonization and local extinction over time. Thus, the curves of these conditions are plotted as scattered “data points”, but not the continuous ones like the others.

For all levels of initial maladaptation, the average fitness of the populations with dormant propagule banks gradually increased, but the fitness dynamics depended on the propagule bank size, initial fitness at the start of the simulations, and reproduction mode (Fig. 3). First, under the condition of initial low fitness of a local population and no or small bank, the local population repeatedly underwent colonization and local extinction, and thus, the curves of these conditions are plotted as scattered “data points,” but not the continuous ones like the others. The local population of colonizing species with low initial fitness remained very low in abundance over the simulation time in the no- or small-bank scenarios (Fig. 2A in Appendix 2). With a larger propagule bank, the fluctuation in fitness and population abundance dynamics weakens. While propagule bank size influences the rate of evolution in both sexual and asexual reproduction scenarios, it does so in opposing ways. The rate of evolutionary adaptation is enhanced by larger dormant propagule banks under sexual reproduction, whereas it is reduced under asexual reproduction.

### Assembly mechanisms and their relative contribution changes with propagule bank size

The relative contributions of ecological processes (i.e., mass effects and species sorting) and eco-evolutionary processes (monopolization at the habitat, patch, or regional scale) to community assembly varied with dispersal rate, propagule bank size, and reproduction mode (Fig. 4). In Fig. 4a if the contribution of a single mechanism was greater than 75%, then the mechanism would be viewed as the main assembly mechanism, i.e., SS, HM, PM, and RM. If none of the single mechanisms contributed more than 75%, then when the ecological processes (i.e., SS+ME) are greater than the eco-evolutionary processes (i.e., HM+PM+RM), the main assembly mechanism would be recognized as an ecological process (denoted as E); otherwise, it would be eco-evolutionary process (denoted as M). In Fig. 4b, the details of the proportions of the contributions of the mechanisms’ contribution are shown.

**Fig. 4.**
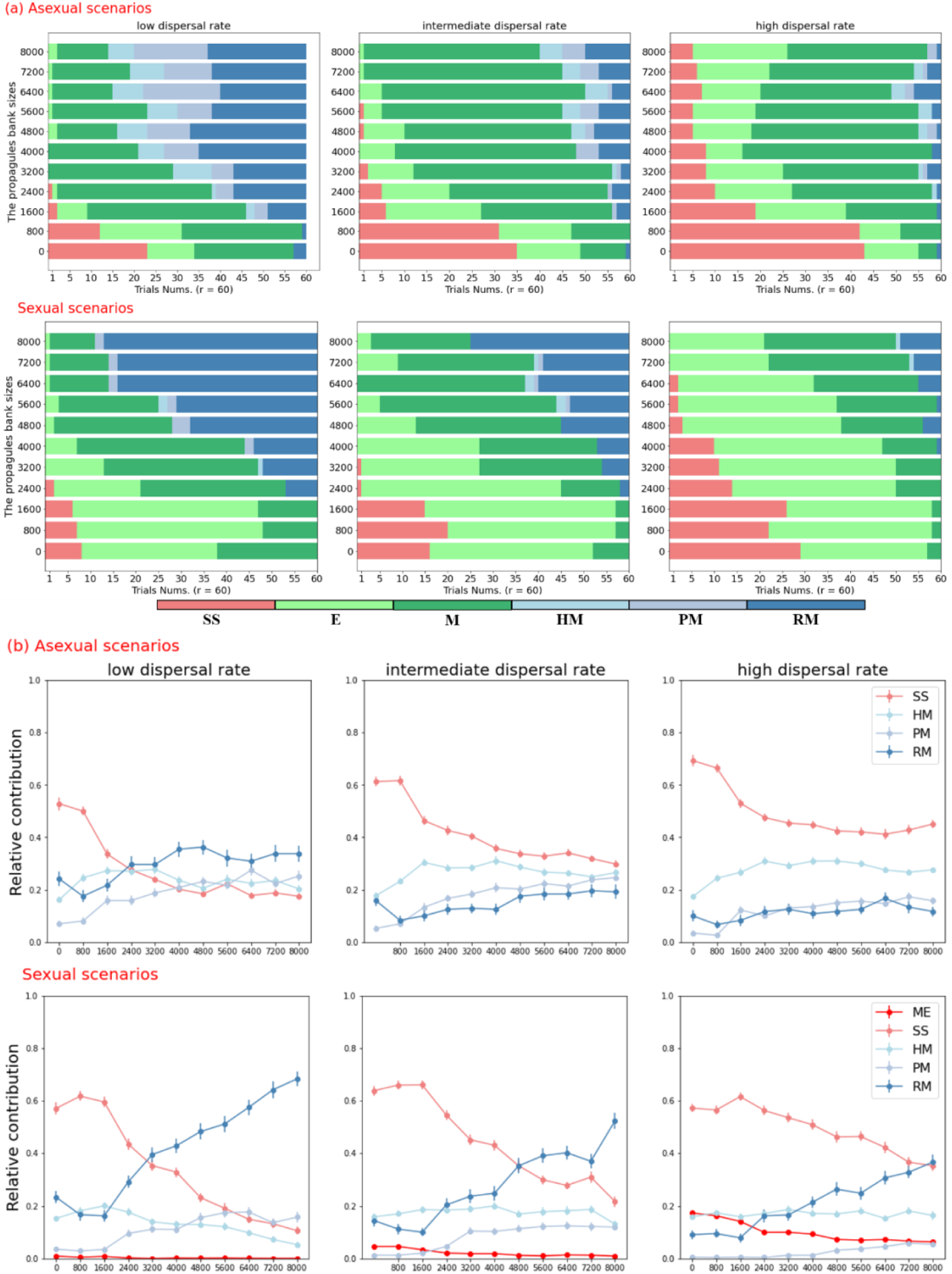
(a) The main assembly mechanisms in the simulation trials in relation to dispersal rate, the propagule bank sizes, and the reproduction mode. Each scenario was simulated with sixty replication trials (*r* = 60) and the variation between trials represents stochastic processes. For a trial, we count the frequencies of individual assembly mechanisms in the metacommunity and the main assembly mechanism is the most prevalent in that trial. Red bars indicate ecological species sorting (SS) with a contribution of over 75% to community assembly; Blue bars with a gradient of darker shades indicate habitat monopolization (HM), patch monopolization (PM), and regional monopolization (RM) with contributions of over 75% to community assembly. If none of a single mechanism contributes greater than 75%, light green bars emphasize that the ecological processes (ME+SS) overtook the evolutionary ones (HM+PM+RM) a bit and the deep green bars emphasize the opposite. **(b)** The relative contribution of each assembly mechanism: mass effect (ME), species sorting (SS), habitat monopolization (HM), patch monopolization (PM), and regional monopolization (RM) in relation to dispersal rate, the propagule pool size and reproduction mode. Error bars are calculated as the standard error of the mean. Note that ‘*propagule bank sizes* = 0’ indicates scenarios without a bank.

In the absence of a dormant propagule bank, community assembly was largely driven by ecological processes, particularly in scenarios of medium to high dispersal. Overall, species sorting was the strongest driver, whereas mass effects were important only in the scenarios of sexual reproduction and high dispersal. From no bank to a small bank, the monopolization effect (RM) became slightly weaker, indicating that the storage effect of the bank protected the inferior species from local extinction in the early stage of their population establishment, when they had to compete with the monopolized species with well-established populations and banks in nearby patches for dispersal. In the presence of a dormant propagule bank, the contribution of species sorting declined with increasing propagule bank size, similar to the mass effect in the sexual scenario. In the scenarios of medium bank size, none of the single assembly mechanisms can become the main assembly mechanism; however, as bank size increases, the eco-evolutionary processes (i.e., HM+PM+RM, denoted as M) gradually overtake the ecological processes (i.e., SS+ME, denoted as E), and finally, the hierarchical monopolization effect overtakes the others. In addition, there was a shift from habitat and patch to regional monopolization in the sexual scenario compared to the asexual scenario.

The hierarchical monopolization effect is strong in scenarios of low dispersal rate, with which a historical contingency of the orders of new arrivals into an empty patch is more important, often leading to a strong priority effect. From low to intermediate and high dispersal, ecological processes, including species sorting and mass effects, became stronger. Higher dispersal with an asexual reproduction mode favors the best pre-adapted species dominating their own niches by the environmental selection process, often contributing to a species sorting effect. Moreover, in scenarios of higher dispersal with sexual reproduction, a source-sink dynamic with micro-evolution makes it possible for an inferior species to persist from local extinction and co-exist with an adapted species, eventually leading to the emergence of a mass effect.

### α-diversity, β-diversity, and γ-diversity

In scenarios without a dormant propagule bank, α-diversity is lower than that in scenarios with a small dormant propagule bank, which implies that the storage effect of the bank favors the α-diversity at the local scale, while the evolution-mediated priority effect in a small bank is still weak. Under the scenario of asexual reproduction, local diversity at the habitat scale was nearly 1.00, indicating that all local habitats were fully occupied by a single species and that mass effects were absent (Fig. 5a). In the sexual scenario, the α-diversity in a single habitat (averaged across the metacommunity) was between 1.00 and 1.20 (Fig. 5g), with α-diversity in some habitats greater than 1.25 (only averages plotted). Such patterns for α-diversity indicate a weak mass effect. The mean α-diversity at the patch or regional scale increased from no bank to small banks, which implies a storage effect, and then decreased with increasing propagule bank size, which indicates an evolution-mediated priority effect (Fig. 5b, c, h, i). β-diversity decreased from no bank to small bank and then increased with larger propagule banks (Fig. 5d, e, j, k). γ-diversity in the global metacommunity fluctuated between 2.0 to 4.0 (Fig. 5f, l) and showed no strong pattern with dormant propagule bank size, indicating that dormant propagule size may reshuffle alpha and beta diversity on a global scale, while reducing alpha diversity in favor of beta diversity as propagule bank size increased in a local scale to regional scale. From a local to regional and global scale, the increasing variation (i.e., the range of the error bar) of alpha to gamma diversity indicates that the stochastic effect is stronger on a large spatial scale. α-diversity at the habitat, patch, and regional scales increased with dispersal rate across the metacommunity except for α-diversity in habitats with asexual scenarios, where α-diversity remains 1.00 (Fig. 5a, b, c, g, h, i). β-diversity decreased at higher dispersal rates in the metacommunity (Fig. 5d, e, j, k).

**Fig. 5.**
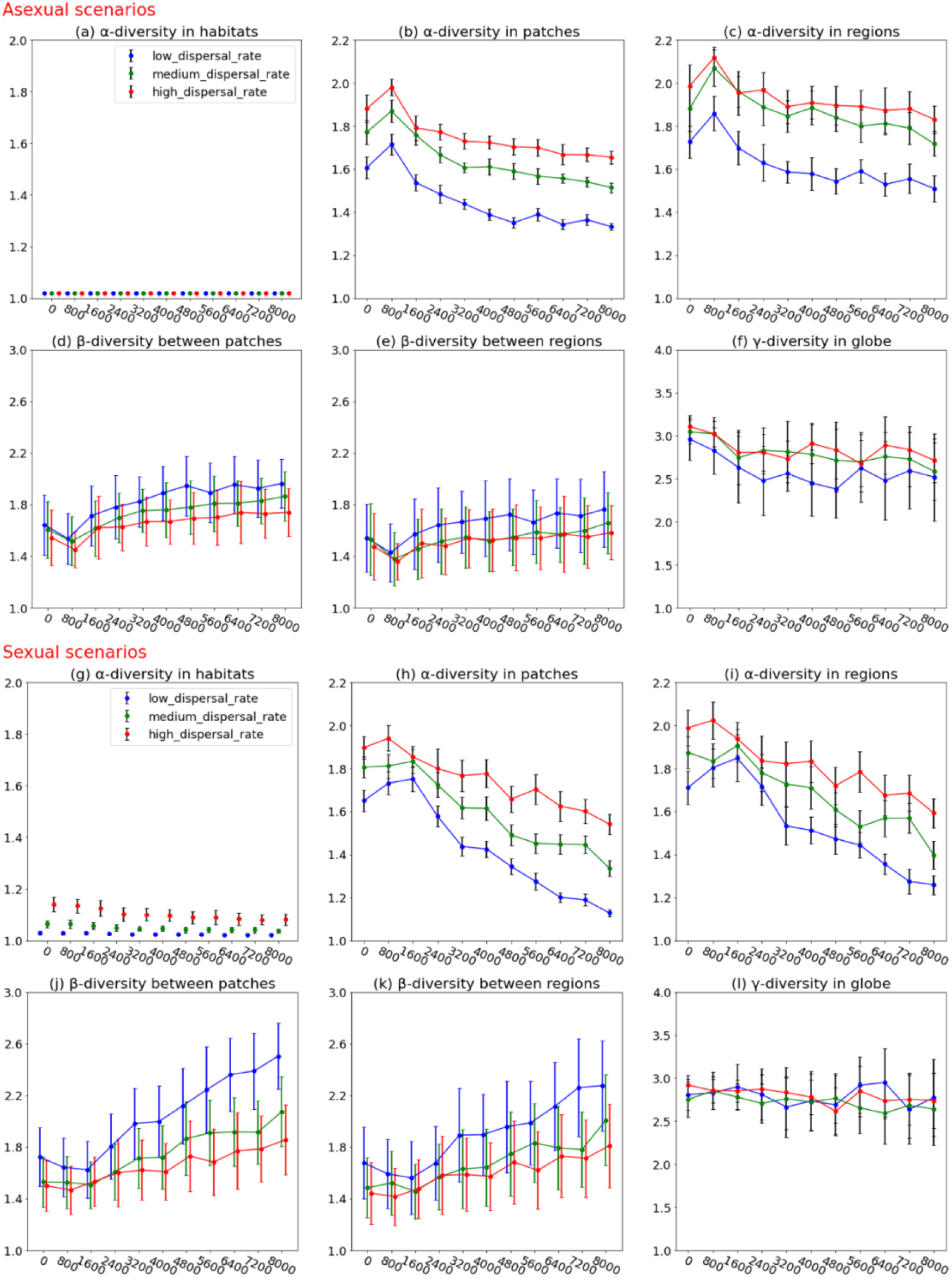
α-diversity in habitats (a, g), patches (b, h), and regions (c, i) are calculated as the average of the inverse Simpson’s index at each spatial scale in the metacommunity in relation to the propagule bank sizes, dispersal scenarios, and reproduction mode. γ-diversity (f, l) is calculated as the inverse Simpson’s index in the global metacommunity. β-diversity (e, f, k, l) is calculated as the ratio of the average of gamma diversity to alpha diversity at patches or regions indicating the variance in species composition between patches or regions. Note that ‘*propagule bank sizes* = 0’ indicates scenarios without a bank.

## Discussion

Dormant propagule banks are expected to strongly affect population and community dynamics (Templeton & Levin 1979, Warner & Chesson 1985, Hedrick 1995) through their impact on both local dynamics and dispersal rates, as well as on metapopulation and metacommunity structures (Wisnoski et al. 2019). Through their influence on both the ecological dynamics and evolutionary trajectories of local populations and the genetic structure in the landscape, dormant propagule banks are also expected to influence eco-evolutionary dynamics in metacommunities in important ways (e.g. De Meester et al 2002, 2016). Yet, the influence of dormant propagule banks on the structure of evolving metacommunities has not been systematically analyzed. Indirect approaches have often been used, whereby bank dynamics are estimated indirectly using aboveground data, assuming a direct link between aboveground populations and underground propagules via dormancy and re-germination processes. For example, a hidden Markov model was built to indirectly estimate the dynamics of underground seed banks for plants. Herein, propagule banks are viewed as a hidden state of the aboveground population (Freville et al. 2013, Borgy et al. 2015, Manna et al. 2017). In the present study, we used individual-based modeling to study the impact of dormant propagule banks on evolving metacommunity dynamics by including propagule dispersal and propagule persistence from an existing metacommunity model (Vanoverbeke et al. 2016). We found that from no bank to a small bank, a storage effect occurred, protecting maladapted species from local extinction and favoring local diversity. We also find that with increasing dormant propagule bank size, (i) in the micro-evolution of a local population, the evolution rate decreases in asexual scenarios and increases in sexual scenarios and the average fitness of the local population over time fluctuated less; (ii) ecological processes such as species sorting and mass effect contribute less to community assembly, while eco-evolutionary processes such as evolution-mediated priority effects become crucial importance; (iii) α-diversity at local scales decreases and β-diversity between local communities increases; Here, we discuss the relative role of propagule bank and propagule dispersal on micro-evolutionary trajectories and population dynamics, on community assembly trajectories, and on the resulting patterns of diversity in the metacommunity.

### The influence of dormant propagule banks on microevolutionary trajectories

Microevolution can occur rapidly in natural communities when sufficient additive genetic variance is present, and populations can persist despite selection (Hendry and Kinnison 1999, 2001, Hairston et al. 1999, Gomulkiewicz & Shaw 2013, Yamamichi et al. 2019, Chaturvedi et al. 2021). Previous studies have shown contradictory effects of propagule banks on the rates of evolution, as they can increase the rate of evolution in the case of fluctuating selection, but can slow down the rate of evolution in the case of directional selection (Hairston and De Stasio Jr 1988, Yamamichi et. Al 2019). In our simulation, we found that in the setting of directional selection, dormant propagule banks slowed evolution in asexual populations and increased the rate of evolution in sexual populations (Fig. 3, Fig. 6a, b, e, f). The overlapping generations of species in the dormant propagule bank and the protection of individuals disfavored by the current selection process can increase generation times, thus maintaining increased genetic variation (Warner and Chesson 1985, Hairston 1996a, Cáceres 1997b). However, in asexual species, this also brings maladapted genotypes back into the population and may thus slow down evolution (Hairston and De Stasio 1988). In sexual species, the contribution of hatching in the old dormant stages to the maintenance of genetic variation for recombination outbalances the genetic load of maladapted genes in the population.

**Fig 6.**
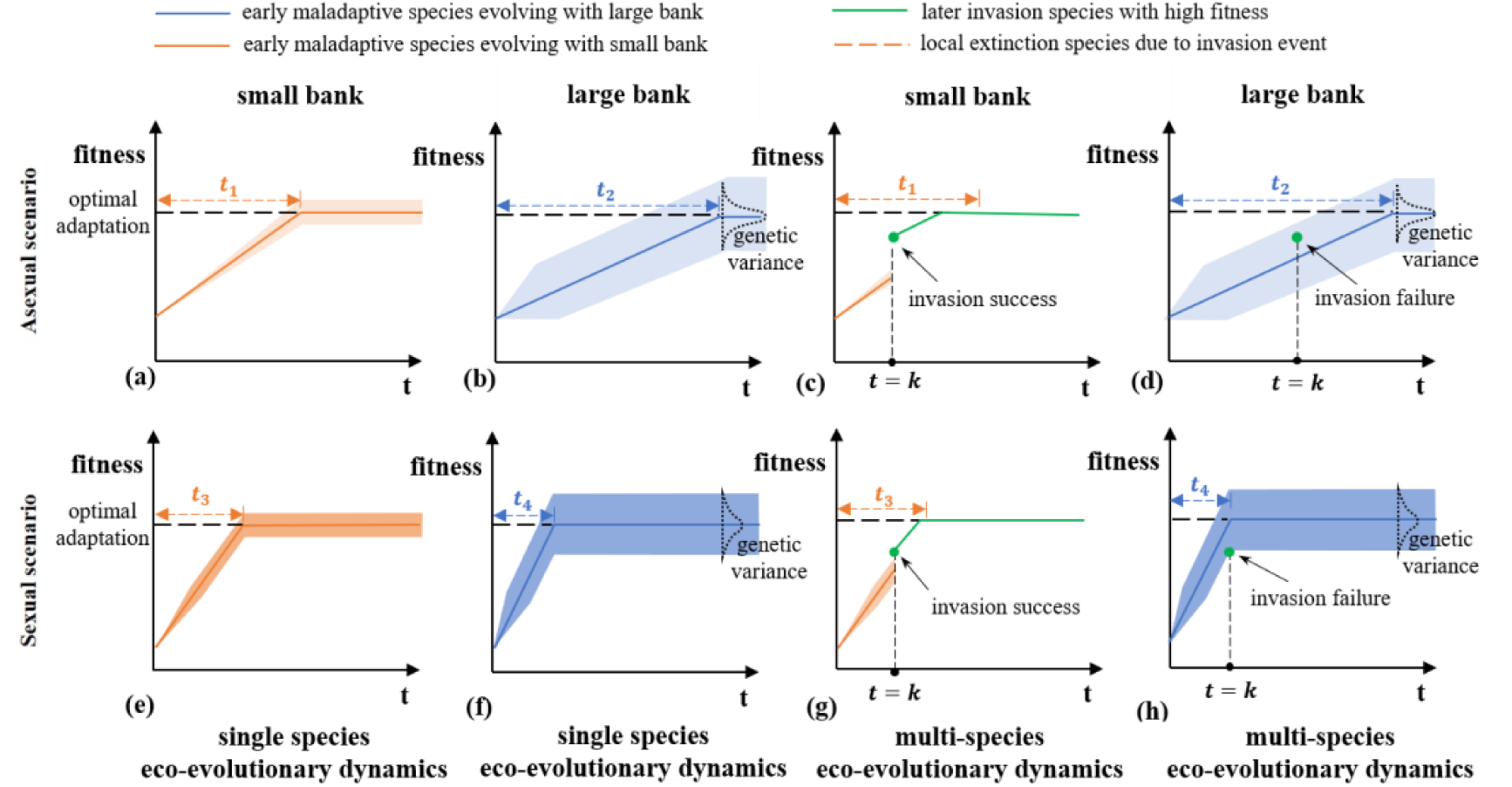
A simplified illustration of the eco-evolutionary dynamics of single species (a, b, e, f) and multi-species (c, d, g, h) in a patch, illustrated by plotting the evolving fitness of initially maladapted species and the buffer effect of the bank, over time in a local patch with either a small or a large dormant propagule bank. We assume that a locally maladapted species evolves improved fitness to the local environment over time t_1_, t_2_, t_3_, t_4_ (depending on the panel). The line labeled with optimal adaptation indicates a local species that would be perfectly adapted to the patch. As the local population is growing, the buffer effect can protect the local population from local extinction owing to fluctuation or invasions. The shading areas indicate an ecological buffer effect due to the growing abundances in the existing population and its dormant propagules, while the depth of the shading colors indicates an evolutionary buffer effect due to the growing genetic variance in the existing population and its dormant propagules. When dispersal among local patches is taken into consideration, colonists with high fitness in the local environment are more likely to outcompete initial resident species during invasion events. If an invasion event point occurs in the zone above shading areas in the plotting, the local population is likely to go extinct, otherwise, the local population can persist. That zone areas can imply the probability of a successful invasion event. We illustrate an invasion event at time-step t=k in the multi-species eco-evolutionary dynamics plots. Solid lines indicate an existing species, while dashed lines show that a species became extinct in a patch. In the asexual scenario (top four panels), (a) it takes a single local maladapted species a shorter time (t_1_) to become adapted with a small bank (orange line) and (b) longer (t_2_) with a large bank (t_2_>t_1_) (blue line). (c) An example of invasion success, in the small banks, a local species is more likely to become extirpated due to population fluctuations and competition with a late arriving species and, (d) An example of invasion failure due to the buffer effect of a bank, a large bank can impede the invasion success of later arriving species due to the ecological buffer effect in the multi-species eco-evolutionary dynamics. In the sexual scenario (lower four panels), (e, f) early maladapted species become adapted more rapidly over time than in the asexual scenario (t_1_>t_3_ and t_2_>t_4_), and consequently early species are less likely to be outcompeted by a later arriving species as the time window for invasion is short. Early species can evolve more quickly in the case of larger dormant propagule sizes (t_3_>t_4_). (g) Compared to the large bank condition, early species are more likely to become extirpated due to a longer period where other species can colonize and outcompete them (t_3_>t_4_) and high population fluctuations in the case of small dormant propagule banks. (h) With a large bank, local maladapted resident species can persist in the patch owing to a shorter period where a competing species could colonize (t_4_) and the buffering effect of a large bank to population fluctuations.

In addition to the contribution of older genetic variants to the population, dormant propagule banks may also support evolutionary rescue owing to their effect on population size. Reznick and Ghalambor (2001) reviewed studies on contemporary microevolution and concluded that the population growth rate following colonization is an important factor in determining the dynamics of the pioneer population. High population growth can reduce the probability of extinction before adaptation can occur (Gomulkiewicz and Holt 1995). We found that a larger propagule bank could stimulate population growth by facilitating the rapid recovery of population size after natural selection, preventing population crashes, and determining the population’s future trajectory. This becomes even more important for maladapted colonists under strong selection, who are liable to crash and become extirpated (Barton & Partridge 2000, Myers and Harms 2011). Thus, in asexual species, a large propagule bank can contribute to the genetic resilience of the local population while also increasing the genetic load from maladaptive variation in a population. In sexual species, where evolution is mainly driven by recombination, dormant propagule banks support adaptation by maintaining high genetic variation and buffering the population in the case of crashes.

### The role of propagule banks in evolving metacommunities

When dispersal and dormancy are taken into consideration, it should be noted that dormant propagule banks not only affect local dynamics (e.g. buffering of population fluctuations, Warner & Chesson 1985) but can also contribute to dispersal. Many species disperse via dormant propagules among suitable patches in a metacommunity, including many plant species that disperse via seeds, fungi, and mosses that disperse via spores, and many zooplankton species that disperse via dormant propagules (Templeton and Levin 1979, Okamura and Freeland 2002, Wisnoski et al. 2019). However, only a few theories combine both dormancy and dispersal processes in a metapopulation framework, let alone in an evolving metacommunity framework, where eco-evolutionary interactions among multiple species create additional complexity (Fréville et al. 2013, Manna et al. 2017, Louvet et al. 2021).

When considering the eco-evolutionary dynamics of multiple species at a local scale, an early maladapted species might fail to persist in the local habitat because of (i) the combination of stochastic effects at small population sizes combined with an incapacity to increase the population size in the harsh environment during the early period of colonization or (ii) competition with better-adapted later-arriving species. The presence of a dormant propagule bank improves the persistence of a maladapted species in a patch in three ways: it enhances survival through dormancy, buffers population fluctuations, and accelerates the rate of evolutionary adaptation in species with sexual reproduction (Fig. 6c, d, g, h). The propagule bank size affects the probability that a dormant individual will successfully survive to the next time step (Buoro and Carlson 2014) and the population density of pioneer species can be better maintained because of the buffer provided by recurrent hatching from the dormant propagule bank (Sarnelle and Knapp 2004). Thus, we can view the propagule bank as a hidden part of the observed population density that represents a larger population size than is appreciated, which can promote a higher per capita growth rate (Dennis 1989). However, for asexual species, a large propagule bank can slow down evolution and thus extend the window of opportunity for colonization by an immigrant species that has a higher fitness than the resident population. A better mutation may have already occurred in the population, while the average fitness is still lower than that of the (one) immigrant (Kilsdonk & De Meester 2021). It seems that a large bank would contain the old genotypes and would protect the new mutant propagules before their germination. For sexual species, dormant propagule banks reduce the time required for adaptation to the local environment by increasing genetic diversity. This increases the probability that the resident population can adapt to the local environment before another species with fitness colonizes the habitat, leading to an evolution-mediated priority effect (De Meester et al 2016).

A large propagule bank usually weakens stochastic events, such as genetic and ecological drift, and weakens the effects of gene flow and dispersal. This is expected to lead to a more deterministic trajectory of microevolution and community assembly once the habitat has been colonized but may increase stochasticity by increasing the importance of historical contingency on an uncolonized or empty habitat, often leading to a strong priority effect. Because of the temporal storage effect, overlapping generations of species are preserved, resulting in the partitioning of the temporal niche (Warner and Chesson 1985, Cáceres 1997b, Chesson 2000b). Therefore, dormancy can act as a buffer against periodically harsh environments that would otherwise lead to local extinction. Such a buffering effect can have a long-term influence on the species community in terms of taxonomic, phylogenetic, and functional diversity (Warner and Chesson 1985, Hairston and Kearns 2002, Lennon and Jones 2011). Similarly, large dormant propagule banks can reduce genetic drift by maintaining genetic variation and increasing the population size (Nunney 2002; Honnay et al. 2008). However, by enhancing the numerical advantage and capacity to genetically adapt to the local habitat, large propagule banks also imply that priority effects may last longer because effective dispersal and gene flow are reduced (De Meester et al. 2002, 2016; Fukami 2005, 2015, Wisnoski et al. 2019).

In the context of eco-evolutionary dynamics involving multiple species in a metacommunity, it is important to include dormant propagule banks as hidden parts of both the regional species pool and local communities. In our simulations, a large dormant propagule bank allowed an early maladapted species to adapt to and monopolize local habitats and, under some conditions, to monopolize multiple habitat patches on a regional scale (i.e., from habitat monopolization to patch monopolization and regional monopolization). At the start of our simulations, all the patches in the metacommunity were empty. Dispersal processes introduce some level of historical contingency in community assembly (Fukami 2015), both in terms of the identity of the species that colonize a given patch and in the timing of patch colonization. As a result, some patches might be colonized early by a species that has the capacity to adapt locally, become locally abundant, and therefore have an increased likelihood of dispersal to neighboring, still empty habitats. This may lead to regional monopolization, a phenomenon that can be reinforced when a dormant propagule bank is involved. All dormant propagule banks were empty at the beginning and accumulated dormant propagules over time. Once a habitat is occupied by a locally adapted population, the resulting dormant propagule bank stabilizes its population abundance over time and enhances the probability of local patch monopolization, as well as the contribution of the local population to the dispersal of propagules in the region, either directly when dormant propagules are dispersal agents or indirectly by supporting larger population sizes.

Moreover, dispersal asymmetries emerge between patches with well-established banks and those with less-established banks. In the assembly of the initially empty metacommunity, owing to historical contingencies, some patches may be colonized and monopolized by a species first, whereas the other patches are still empty or less colonized. Patches with a well-established bank tend to reduce the invasion success of new arrivals, while also providing more emigrants with diverse genetic variation into the other patches. In other words, a patch with a well-established propagule bank makes a more important contribution to the regional species pool than patches with less well-developed propagule banks. Thus, triggered by historical contingencies and intensified by a bank, evolution-mediated priority effects are reinforced by dispersal asymmetries, fostering regional monopolization.

### The effect of dormant propagule banks on species diversity patterns

In the classic metacommunity theory, the focus is on the interaction between regional dispersal and local niche selection (Leibold et al. 2004; Holyoak et al. 2005), but dormancy processes are ignored (Wisnoski et al. 2019). Dormancy can influence species abundance and distribution across spatial and temporal scales (Vandekerkhove et al. 2005, Mahaut et al. 2018, Wisnoski et al. 2019). In our simulations, which included both dormancy and eco-evolutionary interactions, we found that the presence of dormant propagule banks shaped the spatial distribution of species at both the local and regional scales.

At the local scale, propagule banks allow founder species to overcome unfavorable environments until conditions improve (Gioria and Moravcov 2012) and facilitate adaptation to local conditions (De Meester et al. 2002, 2016; simulations in the present study). A larger propagule bank ecologically buffers local population fluctuations, thereby reducing the likelihood of the extinction of founder species. The consequences of the corresponding species diversity, from the absence of a bank to a small bank, are mainly reflected at the local scale. In these cases, the storage effect and the ecological buffer effect will favor the α-diversity in a patch. However, increasing bank size would lead to a loss of species diversity by favoring the evolution-mediated priority effect (see below). Moreover, in the high dispersal and sexual scenarios, the species co-exist in a local habitat due to a mass effect; at first, the maladapted species survive by their source-sink dynamics, and then the microevolution of the maladapted species become adapted to the locality. An increasing bank can reduce effective dispersal and gene flow from immigrants, weakening coexistence of the species by source-sink dynamics.

At a regional scale, as mentioned above, the dispersal asymmetries between patches with well-established banks and those with less-established banks may lead to a regional monopolization effect, which could result in the loss of species diversity. Our spatially explicit model incorporates four spatial scales: habitats within patches, patches in the region, regions, and the globe. A regional monopolization effect can lead to a biodiversity loss among a set of nearby communities, reducing the α-diversity within patches of regions. For distant communities (i.e., local communities among different regions), the increasing size of the dormant propagule bank increases β-diversity among regions. During the early period of colonization, dormant propagule banks allow early propagules to rapidly colonize empty sites. After a colonization event, early-arriving species were more likely to invade adjacent communities. In species where dormancy also increases dispersal (e.g., seeds and dormant propagules that act as dispersal agents), dormancy can facilitate spatial homogenization among adjacent habitats (Wisnoski et al. 2019), leading to regional monopolization and loss of species diversity in a region. Of course, such regional monopolizations are highly context-dependent. Here, we speculate that when a species is specialized in both long-distance dispersal and dormancy and the metacommunity dynamic is somehow isolated from the mainland (i.e., historical contingency due to very limited dispersal from the mainland), a particular state (i.e., local monopolization) is likely to eventually dominate the metacommunity via positive feedback at the local community level (Shurin et al. 2004). Regional monopolization would then occur with a high chance.

## Conclusion

In a metacommunity level, we regard the dormant propagule banks in a metacommunity as a hidden state of a local community, a potential gene pool and a latent part of a regional species pool. The presence of dormant propagule banks and dispersal within metacommunities are important processes that provide mechanisms for species to respond to changing environments, and these processes strongly affect metacommunity structure and dynamics. The presence of dormant propagule banks can affect both evolutionary and ecological processes and thus shape eco-evolutionary dynamics in metacommunities. The banks affect the local microevolutionary trajectories of populations, largely by buffering populations from crashes and maintaining high genetic variation. This, in turn, leads to an increased likelihood of evolution-mediated priority effects that, together with the enhanced likelihood of priority effects driven by higher local population sizes, enhance the historical contingency in determining community composition in local patches. To the extent that dormant stages also enhance dispersal, patch monopolization can lead to regional monopolization. As a result, the presence of dormant propagule banks reduces signals of species sorting and mass effects and enhances the importance of patch and regional monopolization in metacommunities. We show that as a result of these processes, the presence of dormant propagule banks reduces α-diversity within local communities and increases β-diversity between local communities. Collectively, it is important to integrate ecological and evolutionary dynamics, dispersal, and dormancy if one aims to understand how evolving metacommunities assemble or respond to environmental changes.

## Supporting information

An example of video of a simulation

## Acknowledgments

This research was funded by a grant from the NSF of China (grant number 32171538). The High-performance Computing Platform of Jinan University is appreciated for its support in the extensive simulations. LDM was supported by the KU Leuven Research Fund (grant number: C16/23/003).

## Notes

https://doi.org/10.5281/zenodo.13381145

### Competing Interest Statement

The authors have declared no competing interest.

