## Appendix for "The role of dormant propagule banks in shaping eco-evolutionary dynamics of community assembly"

### Appendix 1: Additional details of model designed

*Detail of landscape in the model*

The landscape of the model contains an initially occupied mainland and an initially empty metacommunity. The occupied mainland can provide colonizers to the empty metacommunity.

There exists regions, patches and habitat spatial scales across the metacommunity. There are 4 regions with a regional environment (RE) values gradient of [0.2, 0.4 0.6 0.8] in the metacommunity. In each region, there are 4 same patches. In each patch, there are 4 habitats with a habitat environment (HE) values gradient of [0.2, 0.4 0.6 0.8].

The occupied mainland contains $s$ = 4 species. We consider two traits phenotypes of the species and we define the two traits as *phenotype_1* and *phenotype_2*, which are related to the regional environment (RE) and habitat environment (HE) respectively. The initial genetic traits values of the four species are (0.2, 0.2), (0.4, 0.4), (0.6, 0.6) and (0.8, 0.8), denoted as *(phenotype_1, phenotype_2)*.

At the start of the simulation, the community in the mainland is at equilibrium with the four species are well-adapted to their local habitat in the mainland. Then the species colonize the empty metacommunity from the mainland. The individuals of the four species compete the available microsites in the habitat of the metacommunity and pre-emptive competition is assumed that each microsite can only be occupied by one individual.

*Detail of the resting propagules bank in the metacommunity*

The resting propagules bank is consisted of a set of local banks in the local communities in the metacommunity. We assume that the size of the resting propagules bank is an important factor affecting the strength of the dormancy processes. The size of the resting propagules bank is controlled by a parameter, *size*, which denote the sum of the individual-capacity of all the local banks. The parameter, *size*, is set ranging from 800 to 8000 in the interval of 800. We also set *size=0*, indicating the scenario without a bank. The composition of the resting propagules bank is dynamic over time-steps. All the new-born offspring at current time-step will be all stored in to the bank and some of the offspring in dormancy born at previous time-steps would be removed randomly to keep the number of dormancies in the bank within its maximum (i.e., the parameter, *size*). Moreover, when all the empty microsites in the metacommunity are occupied, the number of new-born offspring in the metacommunity at each time-step is expected to be 800 individuals (see below of details) and they would be put into the bank. Hence, we set the *size* ranging from 800 to 8000 with an 800 interval.

*Detail of species and individual attributes in the model*

Our model is an individual-based model and the attributes of an individual include: *species identifier*, *gender*, *phenotype* and *genotype* information. For *species identifier* (i.e., *sp1, sp2, sp3, sp4*), each individual is labelled as one of the four species identifiers and the label would be inherited to their offspring. For sexual reproduction, only the individuals with the same *species identifier* can mate. As to *gender* of an individual, it will be male or female for sexual scenario and all the individuals will be female for asexual scenario. For phenotype of an individual, two traits of the individual in related to regional environment (RE) and habitat environment (HE) respectively. We call them *phenotype_1* and *phenotype_2* respectively. The genetic-based phenotype is calculated as the mean of genotype plus a random variable with gaussian distribution. The genotype of an individual is a L=40 vector, the first 20 elements of which controls the *phenotype_1* and the second 20 elements of which controls the *phenotype_2*. All the elements for the vector are 0 or 1 to denote the bi-allelic genes.

*Detail of initialization of species and its genetic composition in the mainland*

At *time_step = 0*, four species occupy their optimal habitat of the fours in the mainland. The genotype is created such that the mean of the genotype vector is identical to the initial phenotype values we preset of the species (e.g. if the phenotype is 0.2, we set 16 0s and 4 1s elements randomly in the genotype vector). Then, the phenotype_1 and the phenotype_2 are calculated (mentioned before). During the first 25 time-steps, no dispersal is allowed and the individual go through sexual reproduction to generate standing genetic variation for the population in the mainland by recombination. During the second 25 time-steps, the propagules rain from mainland to the metacommunity is not allowed but the individual can dispersal between habitats in the mainland. Noted that the first 50 time-steps, no matter in asexual scenarios or not, all the species undergo sexual reproduction to create standing genetic variation in the initial population. From the 51 time-steps, the individuals undergo the processes mentioned in the main document.

*Detail of processes in the metacommunity*

The individual in the metacommunity will undergo survival, reproduction & mutation, dispersal and dormancy processes. Pre-emptive competition is assumed that each microsite represent one spatial resource for only one individual. Lottery competition is also assumed that an empty microsites can be re-occupied by individuals randomly chosen from disperses of other regions, other patches or other habitats and the local offspring. For survival process, based on the *phenotype_1* and *phenotype_2* of an individual and the habitat-environment value and region-environment value of a site, we calculated the survival rate of each individual to determine their survival.

Following survival, all the individuals in each habitat will reproduce and the number of new-born offspring is expected to be as follow:

$$I = \sum_{patches} \sum_{habs} b\, N_{f,\ ph}$$

where b is the birth rate related to the reproduction way (i.e., 0.5 for asexual, 1 for sexual) and $N_{f,ph}$ is the number of females in habitat h, patch p, *patches* is the number of patches in the metacommunity, 16 and *habs* is the number of habitats within a patch, 4. Moreover, for asexual reproduction, all the individuals from the habitat are assumed to be female. In addition, when all the microsites in the metacommunity are occupied *I=800* individuals. Mutation process occurs during the reproduction at the same time.

After the reproduction, all the new-born individual will be stored into the bank in priority temporarily. We designed a dynamic propagule bank, the size of the bank over time is expected to be as follow:

$$D_t = I_t + (D_{t-1} - E),$$

where $D_t$, the total number of offsprings in the propagules bank at time step t, is expected to be all the new-born offspring time step t ($I_t$) plus the resting ones inherited from the pool at time step t-1 ($D_{t-1}$) and minus some proportion of offspring in the bank from t-1 time-steps that would be eliminated randomly to keep the total number of offspring in the bank within its carrying capacity. In other word, we note that if $I_t$+$S_{t-1}$ ≤ *the propagules bank size*, then no individuals will be eliminated i.e., E=0.

Later, the migration process takes place and all the individual in the banks play a role as potential colonizers in the metacommunity. All local communities are linked by dispersal of individuals from the local bank to their destination sites. There are three types of dispersal across spatial scales, dispersal within patch, dispersal among patches within region and dispersal across regions. The dispersal strength is controlled by dispersal rates, a parameter, denoted as $m_{across}$, $m_{among}$, and $m_{within}$. The dispersal individual in the metacommunity is expected to be as follow:

$$Disp_{p,\ q} = (I_t + D_t) \cdot m_q$$

where $D_t$ is the total number of offspring within a local propagules bank and $m_q$ is the dispersal rate that depend on the one of the three dispersal pathways where q denotes: *across regions, among patches within regions* or *within patch*.

Further, if the number of dispersal individuals is larger the number of empty sites in the metacommunity, the number of effective established colonizers will be the number of empty sites.

### Appendix 2: details of model under only dormancy process in the exclusion of other metacommunity-levels processes

In this case, we try to understand how the only dormancy process affects the evolution of a maladaptive species in the exclusion of other metacommunity-levels processes. For instance, the gene flow, source-sink dynamics or interspecific competition effect would unpredictably mix up our test for pure dormancy process. Thus, in this scenario, we set dispersal rate equal to 0 and we only consider how the dormancy process affects the population of one single species by setting species num equal to 1.

In the exclusion of the gene flow, dispersal process between local communities is not allowed. In the exclusion of the inter-special interaction, only one initially maladaptive species is provided from mainland by propagules rains. There exist only one habitat-environment and one region-environment in the landscape in a simulation (Fig.1A) and then it tries to get evolutionary adaptation to the environment. The initial phenotype_1 and phenotype_2 of the species is (0.2, 0.2). By varying the habitat-environment and region-environment value at the start of the simulation in this scenario, the initial fitness of the maladaptive species is different (Table.1A). We calculate the mean fitness dynamics of the local population over time and evolution rate to detect the evolution. As results, the evolution of an isolated population is related to the bank size and the initial fitness.

**Figure 1A.** the patchy landscape of the local communities under the propagules' rain of a single non-adapted species. In this case no dispersal among the patchy communities is allowed. The initial mean genotypic trait values (phenotype1) corresponding to the habitat-environment is 0.2 and the initial mean genotypic trait values (phenotype2) corresponding to the region-environment is 0.2. Then, setting different habitat-environment values (no habitat-environment gradients within the patches) and region-environment values (no region-environment gradients within the regions) that the pioneer population is trying to get adapted to via microevolution. Then we detected how the resting propagules bank size affects population evolutionarily and ecologically under different initial selection pressure.

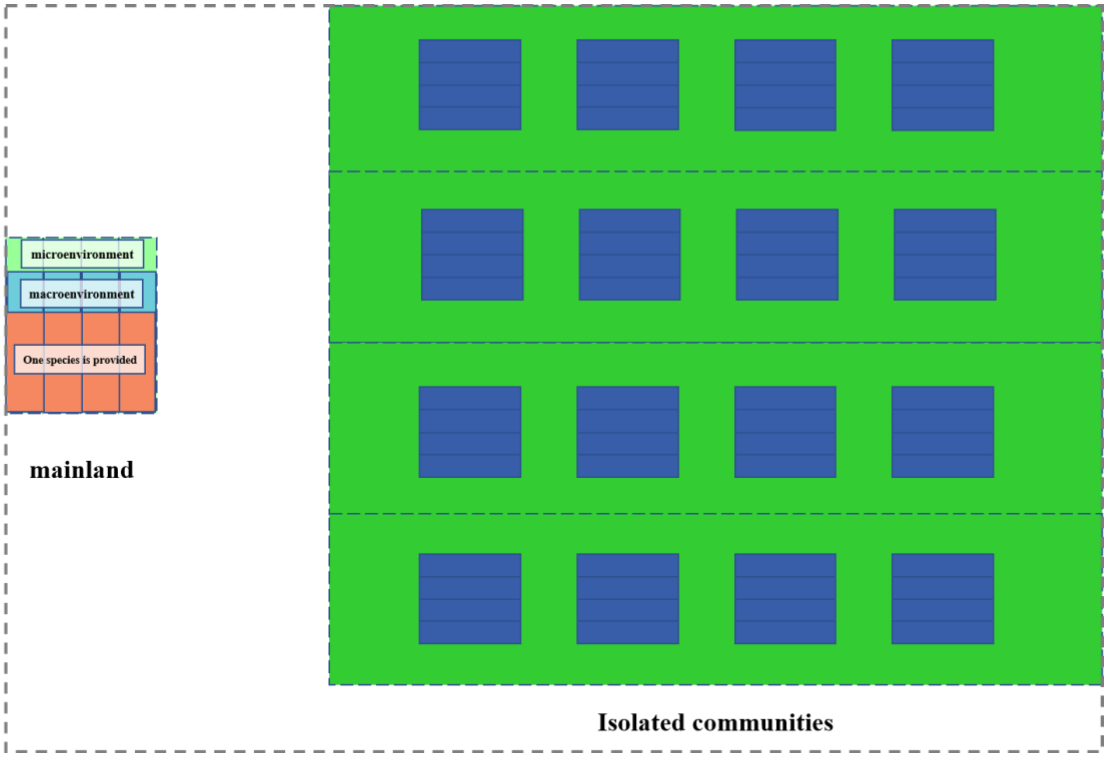

**Table 1A.** for parameters set in the scenario of only dormancy process in the in the exclusion of other metacommunity-levels processes

We detected how the resting propagules bank size affects population evolutionarily and ecologically under different initial fitness. The initial fitness is denoted as the initial mean survival rate of the pioneer species. The initial traits values of the species are fixed (0.2 for phenotype 1; 0.2 for phenotype 2). By varying the habitat-environment values (environment 1) and the region-environment values (environment 2) in each simulation, the population get evolutionary adapted to the environment under high, medium to high, medium to low or low initial selection pressure.

| initial selection pressure | phenotype 1 | environment 1 | phenotype 2 | environment 2 | initial fitness |
|---|---|---|---|---|---|
| **low pressure** | 0.2 | 0.2 | 0.2 | 0.4 | 0.76 |
| **medium to low pressure** | 0.2 | 0.4 | 0.2 | 0.6 | 0.4 |
| **medium to high pressure** | 0.2 | 0.6 | 0.2 | 0.8 | 0.11 |
| **high pressure** | 0.2 | 0.8 | 0.2 | 0.8 | 0.05 |

**Fig 2A.** The total population abundances across the patchy local communities in the exclusion of gene flow and intra-special interaction to estimate the local size effect of the resting propagules banks in the no dispersal case. Within each subgraph, all the non-preadapted colonizers with the same fitness to the local environment at the start of the simulation go through the eco-evolution interaction under the different propagules bank sizes. For the subgraphs from upper to lower, the colonizers population go through the eco-evolutionary interaction under different selected pressure, from the low one, the medium to low one, medium to high one to the high one, which are related to the initial variation between the genetic trait values and the environmental values. The left four panels indicate the population abundances in asexual scenario and the right four panels indicate the population abundances in sexual scenario

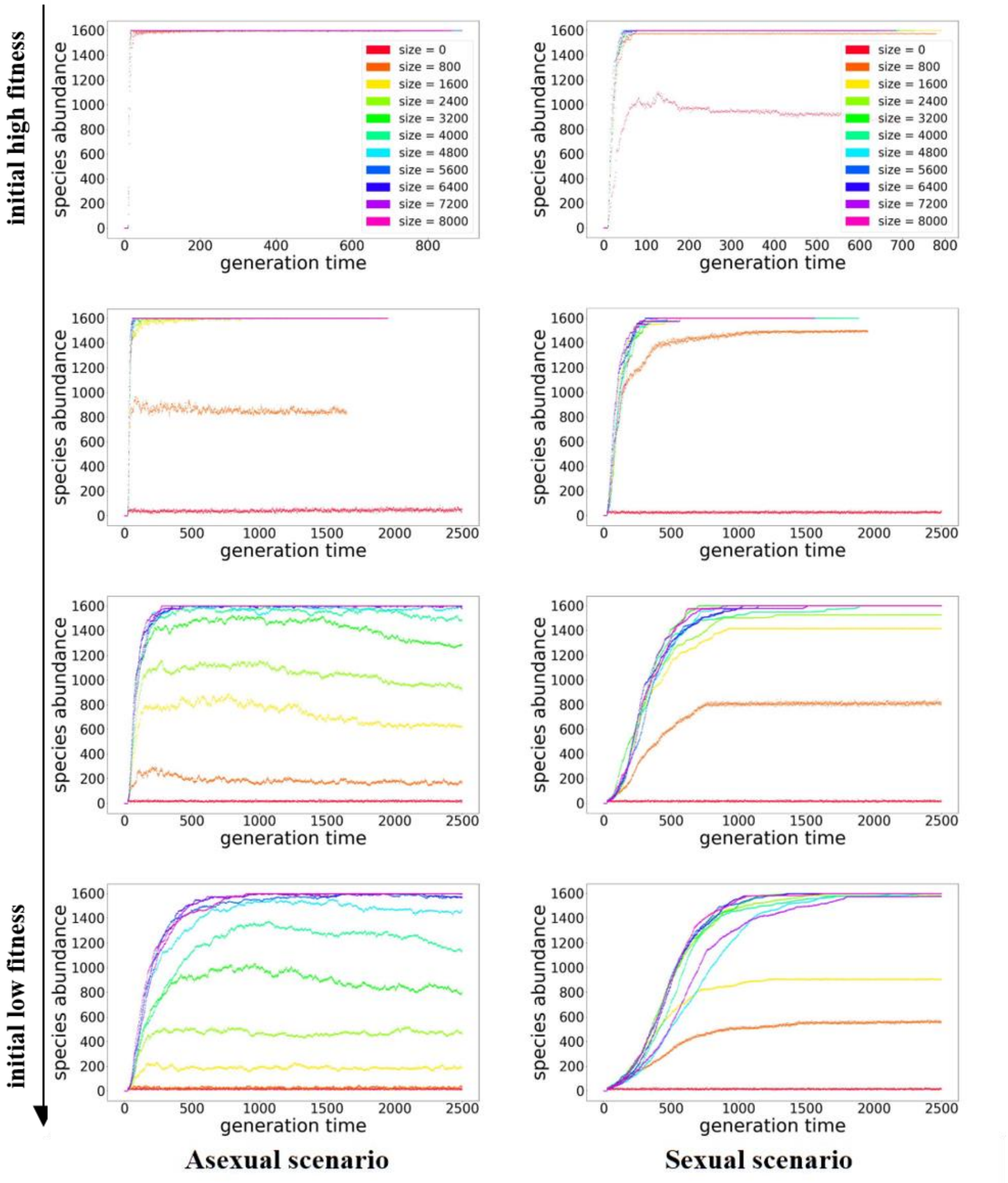

**Fig 3A.** The mean evolutionary rate of the phenotype of the non-adapted population between 10 generations, calculated as the average of 'log of haldanes' between each 10 generations of the population across in the no dispersal case. In the asexual scenarios where evolution driven by mutation is relatively slow, the propagules bank could play a great role in affecting the microevolution of the local non-preadapted species and in the sexual scenario where evolution is main driven by both mutation and recombination, the propagules bank take a less credit for the evolution of the local species. The values of 'log10 of haldanes' under low selection pressure, medium to low selection pressure, medium to high selection pressure and high selection pressure are denoted as blue, red, green and purple respectively.

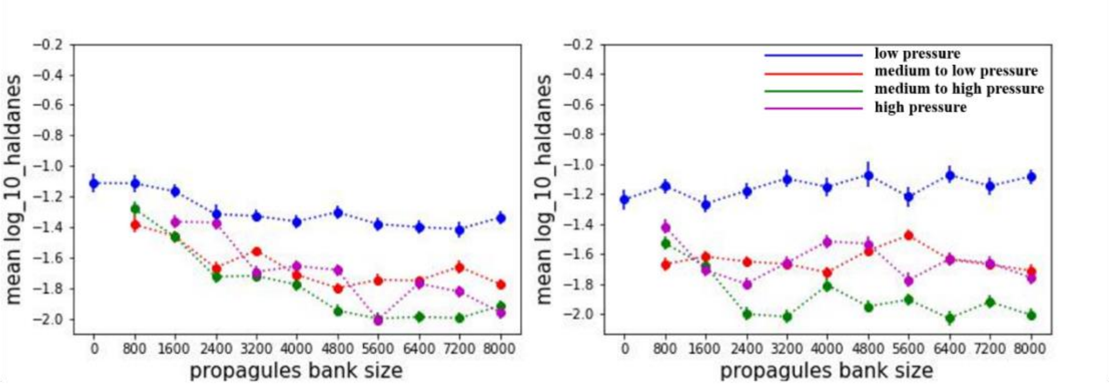


**Asexual scenario** **Sexual scenario**

We estimated the rate of adaptation of a population to the local habitat as the log change in trait values scaled by the pooled standard variation per generation (*haldanes*)

$$\text{haldanes} = \frac{\left(\frac{x_{n+1}}{S_p}\right) - \left(\frac{x_n}{S_p}\right)}{g}$$

where $x_n$ is the average trait at *generation n, g* is the number of generations, and $S_p$ is the pooled standard deviation of the trait across generations ($\sqrt{[\mathrm{SS}_1 + \mathrm{SS}_2]/[(\mathrm{n}_1 - 1) + (\mathrm{n}_2 - 1)]}$), where $SS_1$ or $SS_2$ is the standard deviation of trait values in the population at *generation n* or *generation n*+1, and $n_1$ or $n_2$ is the population size at *generation n* or *generation n*+1.

#### Appendix 3: details of the calculation of the generation time

The *generation time* of the population is calculated as (Gotelli 2008):

$$G = \frac{\sum l_x b_x x}{\sum l_x b_x} \ldots\ldots\ldots\ldots \quad (1)$$

where $l_x$ denotes the survivorship of the population, $b_x$ denotes the fecundity of the population and $x$ denotes the time-steps in the simulation model. With each microsite holding only one individual, the number of individuals of a population in the local community will be limited at the maximum capacity of the local habitat during the its evolution and adaption in the simulation. With the limitation, the population can stay in a relevant stability of the its size in the simulation and we assumed that

$$R_0 = \sum l_x b_x = 1 \cdots\cdots\cdots\cdots \quad (2)$$

The survivorship of the population at each time step can be calculated as:

$$l_x = S^x \cdots\cdots\cdots\cdots \quad (3)$$

Given formular (1), (2) and (3), we can deduce that

$$\sum l_x b_x = b_x \sum_{x=1}^{\infty} s^x = b_x \cdot \lim_{x \to \infty} \frac{s - s^x}{1 - s} = b_x \cdot \frac{s}{1 - s} = 1$$

Then,

$$b_x = \frac{1 - s}{s} \cdots\cdots\cdots\cdots \quad (4)$$

We can calculate the generation time of the population, given formular (1), (2), (3) and (4)

$$Generation\ time = \frac{\sum l_x b_x x}{\sum l_x b_x} = b_x \sum_{x=1}^{\infty} (x \cdot s^x) = 1 / (1 - S) \cdots\cdots\cdots\cdots \quad (5)$$

Hence, the generation time is related to the mean survival rate of population. The generation time of the population can be change over time and we approximately calculate the generation time dynamics according to formular (5) using the data of survival rate over time during the simulation.

### Appendix 4: the work flow of the model and programming

We input the parameters and values at the start of the simulation and then we initialize the metacommunity and the species in the mainland. Individual in the model undergoes nature selection, reproduction, mutation, dormancy and dispersal modules for 5000 time-steps to reach the equilibrium and finally we output and visualize the result.

**Fig 4A. the work flow of the model and programming**

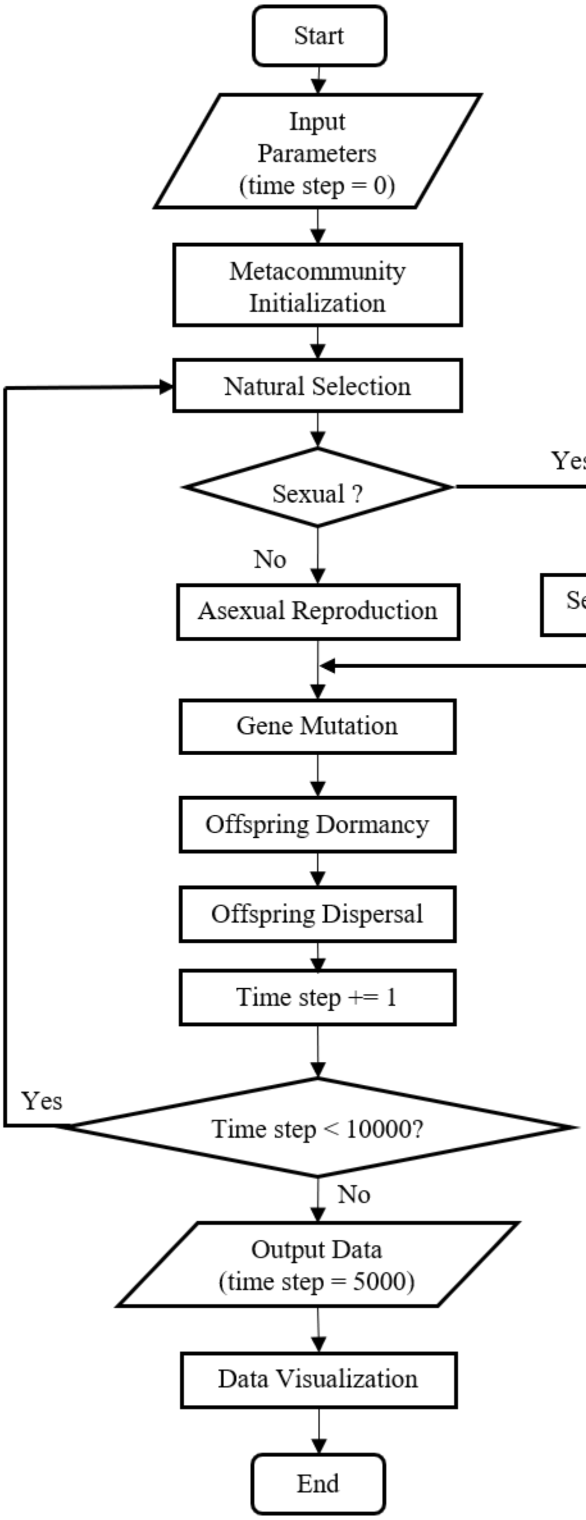

### Appendix 5: Decision making table for each assembly mechanisms

**Table 2** Decision making table for each assembly mechanisms associated with species diversity at habitat, patch, region, global scales and species traits (Vanoverbeke et al. 2016)

| | Mass effect | Species sorting | Habitat monopolization | Patch monopolization | Region monopolization | Global monopolization |
|---|---|---|---|---|---|---|
| **Habitat diversity** | **≥ c** | **< c** | **< c** | **< c** | **< c** | **< c** |
| **Species trait** | | **Pre-adapted** | **Non-preadapted** | **Non-preadapted** | **Non-preadapted** | **Non-preadapted** |
| **Patch diversity** | | | | **< c** | **< c** | **< c** |
| **Region diversity** | | | | | **< c** | **< c** |
| **Global diversity** | | | | | | **< c** |

Denote: the value of c = 1.25 is calculated such that a dominant species has at least a portion of 90% in the case

### Appendix 6: Example movie of a simulation showing community assembly dynamic in the metacommunity over time.

The configuration of the movies consists of 17 patches, with the mainland denoted as *'patch 0'* and the 16 patches in the metacommunity denoted as *'patch1'* to *'patch16'* respectively. Four colors in the movie are precented (black, purple, red and orange) to describe the species identifier that each individual is belongs to.

### Appendix 7: Parameters and values used in the model

**Table 3S** Parameters and values used in the model

| Parameter | Description | Value(s) or Mean value(s) |
|---|---|---|
| **Environment** | | |
| *regs* | Number of regions | 4 |
| *patches* | Number of patches within a region | 4 |
| *habs* | Number of different habitat types | 4 |
| *RE* | Regional environment with metacommunity-level variation | 0.2 to 0.8 in step of 0.2 |
| *HE* | Habitat environment with patch-level variation | 0.2 to 0.8 in step of 0.2 |
| $\sigma_e$ | Microsite micro-environmental variation | 0.025 |
| $RE_m$ | RE-related micro-environmental value in a microsite | mean (RE)+gaussian(0,$\sigma_e$) |
| $HE_m$ | HE-related micro-environmental value in a microsite | mean (HE)+gaussian(0,$\sigma_e$) |
| *n* | Number of microsites per patch | 100 |
| **Demography** | | |
| *d* | Baseline mortality | 0.1 |
| *b* | birth rate per time-step per female | 0.5 (asexual) or 1 (sexual) |
| $\omega$ | Width of the survival rate function | 0.5 |
| $\sigma_z$ | Non-genetic phenotype variation of an individual | 0.025 |
| $\sigma_g$ | Standing genetic variation of individual phenotypes in the mainland at the start of a simulation | 0.045 |
| *G* | Genotype of an individual | a vector with a length of *L* |
| $RE_i$ | RE-related phenotype value of a specific individual | mean(G)+gaussian(0,$\sigma_z$) |
| $HE_i$ | HE-related phenotype value of a specific individual | mean(G)+gaussian(0,$\sigma_z$) |
| ***Disp. scenarios*** | | |
| $m_{across,}$ | Long-distance dispersal rate across regions | $m_{across} \leqslant 0.1$ |
| $m_{among,}$ | Medium-distance dispersal rate among patches in a region | $m_{among} \leqslant 0.1$ |
| $m_{within}$ | Short-distance dispersal rate between habitats in a patch | $m_{within} \leqslant 0.1$ |
| low disp. rate | $m_{across} = 0.001$, $m_{among} = 0.005$, $m_{within} = 0.01$ | $m_{across} < m_{among} < m_{within}$ |
| medium disp. rate | $m_{across} = 0.005$, $m_{among} = 0.01$, $m_{within} = 0.05$ | $m_{across} < m_{among} < m_{within}$ |
| high disp. rate | $m_{across} = 0.01$, $m_{among} = 0.05$, $m_{within} = 0.1$ | $m_{across} < m_{among} < m_{within}$ |
| **Colonization** | | |
| *c* | Colonization rate from mainland | $c = m_{across}$ |
| **Genetic** | | |
| *L* | Number of genes controlling two phenotypes | 20 |
| $\mu$ | Mutation rate | $10^{-4}$ per gene per time-step |
| **Model** | | |
| *time_step* | Total time-steps that the model runs | 5000 |
| *r* | Replication runs | 60 |
| *size* | The individual carrying capacity of the propagule bank | 0, 800, 1600, …, 8000 |
